# Development and Characterisation of Equine Gastric Organoids

**DOI:** 10.1101/2025.11.28.690112

**Authors:** Tanith Harte, Helen Todd, Stewart T.G. Burgess, Jo Moore, Beth Wells, David Smith

**Affiliations:** Moredun Research Institute, Pentlands Science Park, Bush Loan, Penicuik, Midlothian EH26 0PZ

**Keywords:** Horse, organoid, 3D cell culture, stem cell, equine organoid, equine gastric organoid, equine gastric mucosa, organoid differentiation

## Abstract

The healthy equine stomach is of essential importance to equine digestive health and overall wellbeing. Diseases of the gastric mucosa such as gastric ulcers are highly prevalent in equids, particularly amongst racing thoroughbreds, and present a significant challenge to animal health and welfare. Despite this, there is currently no lab-based model capable of accurately representing the cellular diversity and cellular interactions within the equine gastric mucosa to aid in development of treatment options. We present here the first report of equine gastric organoids derived from glandular mucosa tissue. Through bulk RNA-sequencing and immunofluorescence we show these gastric organoids are representative of the tissue whilst retaining the individual gene-expression properties of the animal they were derived from. We also show these organoids are stable over multiple passages and can be differentiated towards different cell-types to upregulate protein and gene expression of specialised cell markers such as gastric foveolar cell associated mucin 5ac, *gastrokine 1 & 2* and *trefoil factor 2*. This model establishes the first physiologically relevant *in vitro* model of the equine gastric epithelium, representing a significant opportunity for developing understanding and therapeutic treatments of gastric diseases in equids.

## Background

The equine stomach, while relatively small in comparison to the rest of the equine gastrointestinal (GI) tract, plays a critical role in equine gastrointestinal health (Krunkosky et al., 2017) as horses are grazing animals, therefore their glandular stomach constantly secretes acid to facilitate digestion of foraged food (Campbell-Thompson & Merritt, 1987). This glandular mucosa is lined with a columnar epithelium composed of secretory and absorptive epithelial cells that form distinct glandular structures (T.S Mair, 2017), which can be further divided into sub-regions where specialized epithelial cells are found (Figure 1). The differentiation of epithelial stem cells into these specialized cells is determined by a gradient of different signaling molecules, produced by cells of the underlying submucosa.

**Figure 1:**
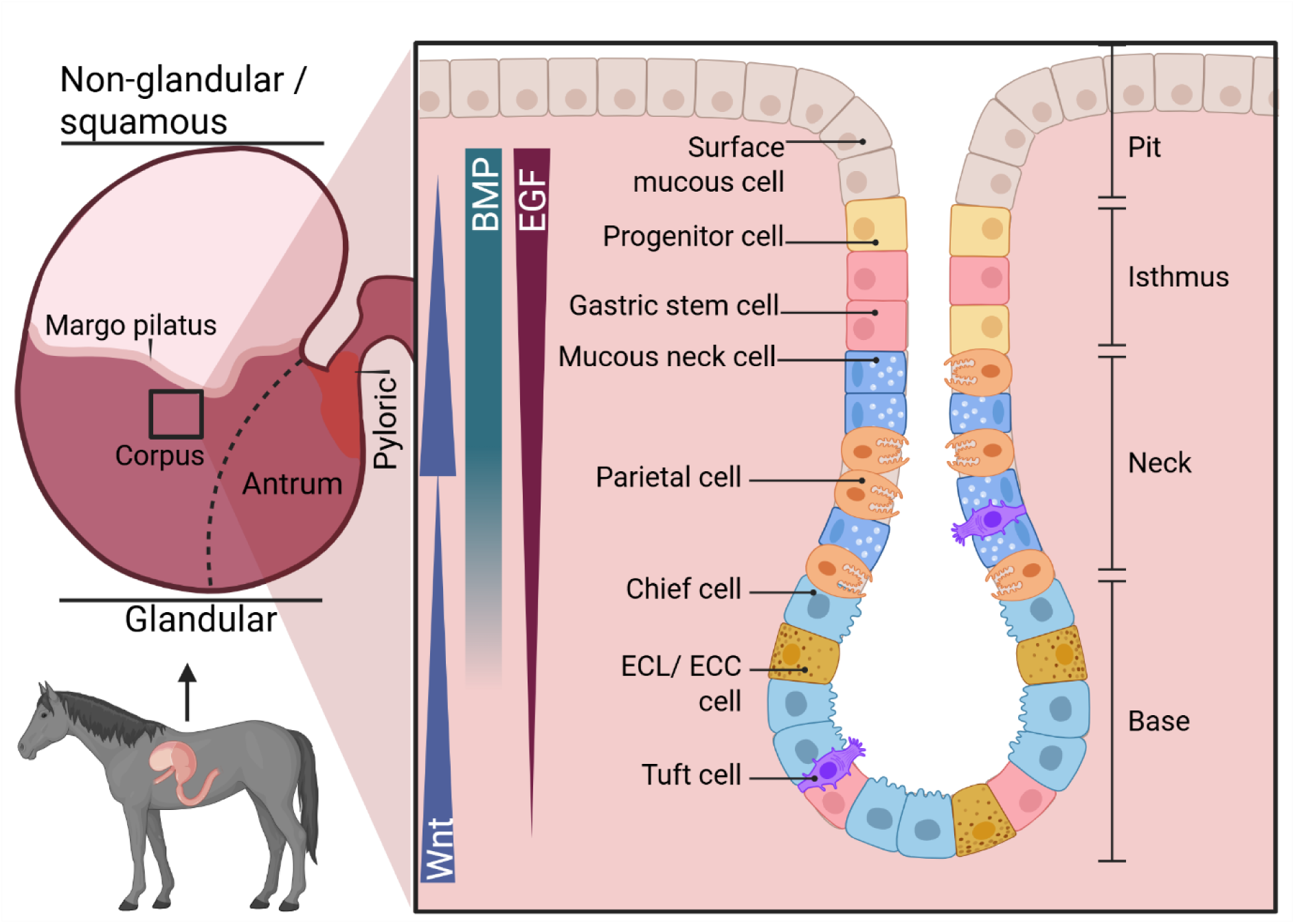
Cellular diversity of the equine glandular mucosa. The equine stomach is divided into 2 sections, the squamous or non-glandular mucosa which has a thick keratinized epithelial surface layer, and the glandular mucosa which consists of columnar epithelial cells which line the gastric glands and which produce acid and hormones to facilitate digestion. The glandular and non-glandular regions are separated by the margo plicatus. The pyloric sphincter separates the stomach from the small intestine (duodenum). Inset shows a fundic gland containing diverse cell types divided by distinct regions within the gland. The pit region forms the uppermost layer of the gland and contains surface mucous cells (or foveolar cells) which express gastric mucus. In the isthmus, gastric stem cells in varying states of differentiation are found. In the neck of a fundic gland, acid-producing parietal cells, tuft cells and hormone producing enteroendocrine cells (EECs) or enteroendocrine-like cells (ECL) form the largest area of the gland. At the base, pepsinogen-producing chief cells are predominantly expressed, though tuft cells and EEC/ECLs can also be found here. Differentiation of the gastric stem cells in either direction is controlled by a gradient of molecules expressed by surrounding mesenchymal cells, including epidermal growth factor (EGF), Bone-morphogenic protein (BMP) and Wnt produced by surrounding mesenchymal cells. Image created with Biorender, with reference to (Hong et al., 2025) and (Krunkosky et al., 2017).

Understanding the cellular diversity and mechanics of the equine stomach is essential for advancing knowledge in areas such as veterinary gastroenterology, equine nutrition and welfare, particularly given the predisposition of equids to ailments such as gastric ulcers, which are thought to result from the production of acid from the glandular region (Vokes et al., 2023).

To date, there are very few suitable lab-based alternatives for research into equine diseases, with research relying on flat monocultures consisting of a single cell type, or the use of live animals. In 2022, the UK Home Office reported the use of horses in approximately 11,000 experimental procedures (Home Office, 2024), despite their assignment as a companion animal. The use of live animals is not only time-consuming and expensive but also comes with considerable ethical concerns. While a number of cell lines have been established for equine research (John et al., 2000; Oguma et al., 2013; Rooney et al., 2023) these fail to capture the diversity of cells within complex GI tissues, diverse signaling between different cell types and lack the cohesive structure. There is, therefore, a need to develop a lab-based model that better recapitulates the gastric mucosa, which is better suited to investigate mechanisms of equine nutrition and host-pathogen interactions, towards advancing knowledge on equine gastric diseases and further improve equine welfare.

In recent years, research into gastroenterology has advanced significantly due to the development of GI-tract organoids. Organoids are *in vitro* cultured microtissues capable of self-organising into 3D structures, which more accurately represent the diversity of cells and the structure of these cells present in the target host tissue. Organoids can be developed either from induced pluripotent stem cells (iPSCs), embryonic stem cells or from host-derived stem cells, such as those found in the crypts and glands of the GI-tract. Tissue derived organoids are particularly effective at recapitulating the target tissue and have been shown to retain signatures associated with the individual animal from which they are derived (Faber et al., 2022). To date, organoids have been used as tools to investigate the interaction of pathogens (Faber et al., 2022; Hofer et al., 2025) in the stomach, the development of gastric ulcers (Xu et al., 2022) and gastric cancer and drug development in humans (Kan et al., 2025). All of these diseases are applicable to equine research (Figure 2), meaning a similar model for the equine stomach would be invaluable. Furthermore, while there are reasonably successful pharmaceuticals available to treat gastric ulcers, there is limited understanding of the mechanisms and interaction with the equine stomach. Despite the extensive potential for applications, a model has not yet been developed for gastric organoids derived from horses.

**Figure 2:**
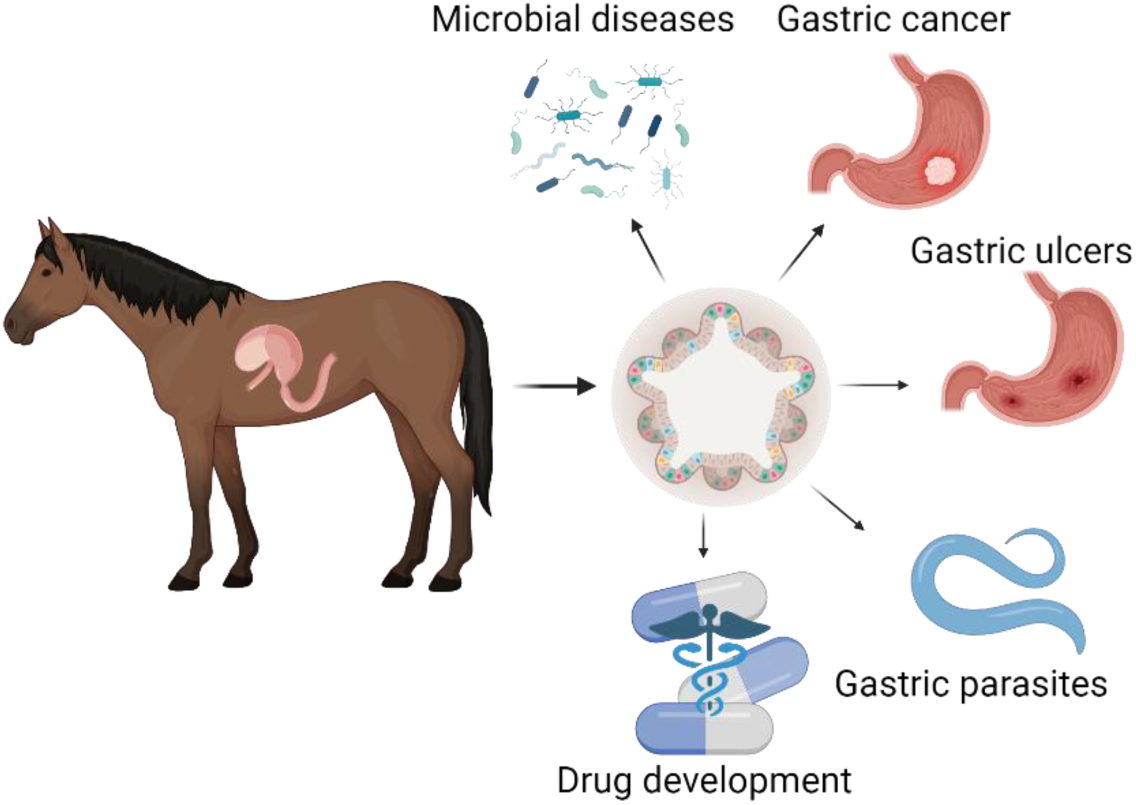
Potential applications of equine gastric organoids. Equine gastric organoids could be used to investigate a variety of diseases affecting horses such as viruses, bacterial and parasitic infections, development of gastric ulcers and formulation of drug toxicity and development.

Here we show methods for deriving organoids from equine glandular gastric tissue, methods for differentiating these organoids to more accurately represent the pit region of a gastric gland and show that these organoids retain similar RNA expression patterns to the original host tissue in terms of gene expression over multiple passages.

## Materials & Methods

### Animals

This study was performed under MRI Equine Grass Sickness Biobank AWERB approval 160321, in line with 3R principles: primarily the reduction of animals for use in scientific research and the reduction of animals used for research. All gastric tissue used for this study was collected from animals being euthanized for welfare reasons unrelated to scientific experimentation. Animals sampled included a variety of ages, sex and breeds. All animals were euthanized by a licensed veterinarian who also performed the post-mortem and tissue collection.

**Table 1:** Metadata for horses used in this study. Samples were collected opportunistically, resulting in significant variation in age and breed.

| Horse ID | Sex | Age | Breed |
| --- | --- | --- | --- |
| 1 | Gelding | 10 | Shetland Pony |
| 2 | Gelding | 4 | Shetland Pony |
| 3 | Mare | 3 | Shetland Pony |
| 4 | Gelding | 15 | Fell pony |
| 5 | Mare | 22 | Thoroughbred |
| 6 | Mare | 20 | Highland Pony |

### Tissue collection

Gastric tissues were collected at post-mortem, similar to the approach used in Smith *et al*. (Smith et al., 2021).

Briefly, approximately 10 cm x 10 cm sections of gastric tissue from the fundic gland region of the equine stomach were collected, briefly rinsed in clean water and then placed in wash media (sterile, ice-cold HBSS with calcium and magnesium(Gibco, 11560616) containing 25 μg/ml gentamycin (Fisher Scientific, 11530506), 100 U/ml penicillin/streptomycin,100 μg/ml Normocin® (InvivoGen, Antnr-05) and 2.5 μg/ml Amphotericin B (Fisher Scientific, 11526481). Samples were stored in wash media on ice until processing (within 3 hours of collection).

### Glandular gastric organoid generation

Tissue samples were rinsed 3 times using fresh wash media then placed onto a sterile petri dish. The mucus layer was removed by gently scraping the tissue with a sterile glass slide and discarded. The epithelial layer was then harvested by scraping again with a clean glass slide using firm pressure and transferred to a Falcon tube containing fresh wash media. Samples were centrifuged at 400 x *g* for 2 minutes. The supernatant and the top layer of the pellet containing a secreted mucus layer was removed and discarded. This process of centrifugation and mucus removal was repeated until no visible mucus layer remained and the wash media was clear.

Epithelial scrapings were then placed into 25 ml digestion medium (25 ml DMEM (Gibco, 13345364), 25 μg/ml gentamycin, 100U/ml penicillin/streptomycin, 100 μg/ml Normocin®, 50 U/ml collagenase (Merck, C2674), 10 U/ml dispase (Roche, 4942086001)) and incubated in a shaking incubator at 80 rpm for 40-45 minutes at 37 °C. Following digestion, samples were allowed to settle at room temperature (RT) for approximately 2 minutes to allow large, undigested tissue sections to sink. The supernatant containing gastric glands was transferred to a sterile 50 ml Falcon tube and gland integrity and quantity assessed via inspecting 10 µl of supernatant by light microscopy. Samples were then centrifuged at 400 x *g* at RT for 2 minutes and subsequently washed with 30 ml of wash media. Samples were then centrifuged for 5 minutes at 400 x *g*, and crypts resuspended in seeding media (Advanced DMEM/ F12 (Gibco, 11540446), 1x B-27 minus vitamin A (Fisher Scientific, 12587010), 1x N-2 (Fisher Scientific, 11520536), and 100 μg/ml Normocin®) at approximately 500-1000 crypts per 100 μl of media.

### Organoid culture

Approximately 1,000 gastric glands from glandular tissue were combined with 100 µl seeding media for each animal. Re-suspended glands (100 μl) were combined with 150 μl of BD Growth Factor-reduced Matrigel® matrix (4 °C)(Fisher Scientific 11543620), and 50 μl was pipetted directly into the center of consecutive wells of a preheated (37 °C) 24-well tissue culture plate to create domes. Culture plates were then incubated at 37 °C, 5% CO_2_ for 45 minutes to allow Matrigel® polymerization. Organoid growth media (OGM; Complete Intesticult™ Growth Medium (mouse) containing 500 nM Y-27632 (Tocris Biotechne, 1254), 10 μM LY2157299 (Cambridge Bioscience, CAY15312), 10 μM SB202190 (Cambridge Bioscience, CAY10010399), 1x B-27 minus vitamin A, 1x N-2, 50 μg/ml gentamycin and 100 μg/ml Normocin®) was prewarmed to 37°C. OGM (550 μl) was then added to each well. Organoids were incubated at 37 °C, 5% CO_2_ and media completely replaced every 2-3 days. Organoids were cultured for approximately 5-7 days prior to passage.

### Passaging glandular gastric organoids

Organoid growth media (OGM) was removed from cultured organoids and discarded. The Matrigel® matrix was dissolved by adding 1 ml ice-cold advanced DMEM/F12 to the well and rapidly pipetting up and down 5-6 times. The suspension was then moved to a 15 ml falcon tube. Each well was rinsed a second time with ice-cold advanced DMEM F/12 and pooled into the same 15 ml sample Falcon tube containing the suspended organoids. Samples were centrifuged at 200 x *g* at RT for 5 minutes to pellet organoids, and supernatant removed and discarded. Glandular gastric organoids (GL-organoids) were then resuspended in 200 µl of seed media and mechanically disrupted by repeated pipetting (approximately 30-40 times) using a 200 μl pipette tip bent at a 90° angle. Organoid fragmentation was confirmed by observing a 10 μl drop under a light microscope, then 100 μl of fragments combined with 150 μl Matrigel® and placed into a 24-well tissue culture plate as described above.

### Organoid differentiation

Organoids were differentiated at P6-7. After passaging, GL-organoids were fed standard OGM for 3-4 days to allow recovery from passage and sufficient growth and branching as to allow for differentiation, estimated visually by an approximately 3-fold expansion in size. After 3-4 days, OGM was removed and replaced with 550 µl of pre-warmed organoid differentiation media (ODM; complete Intesticult™ Differentiation Medium (human) (Stemcell Technologies, 0600), 500 nM Y-27632, 1x N-2 Supplement, 1x B-27 Supplement, 50 µg/ml gentamycin, 100 μg/ml Normocin®) and incubated at 37 °C, 5% CO_2_ for a further 3-4 days.

### Organoid and gland cryopreservation

Surplus glands that were not immediately seeded into Matrigel® were cryopreserved by centrifuging at 400 x *g* for 2 minutes, removing the supernatant and resuspending in Cryostor® CS10 (Stemcell Technologies, 100-1061)at approximately 500 glands or 100,000 cells per milliliter. This suspension was then transferred to a cryovial and stored in a cryogenic freezing container overnight at −70 °C, before being transferred to −150 °C for long-term storage. Organoids at different passages were cryopreserved by gently releasing the organoids from Matrigel® domes using 1 ml cold (~4 °C) Advanced DMEM/F12, transferring to a 15 ml Falcon tube and bringing up to 10 ml with additional Advanced DMEM/F12, then centrifuging at 300 x *g* for 2 minutes at room temperature. The supernatant was removed and organoids resuspended in chilled Cryostor® CS10 (at ~500 whole organoids per cryovial). Cryovials were immediately placed in a cryogenic freezing container and frozen overnight at −70 °C before moving to −150 °C for long-term storage.

Cryopreserved organoids or glands were resuscitated by rapidly thawing cryovials in a water bath at 37 °C, then quickly transferring into a 15 ml Falcon tube containing 8 ml of Advanced DMEM/F12. Samples were pelleted by centrifugation at 400 *x g* for 5 minutes at 4 °C, then resuspended in seeding media. The suspended, resuscitated organoids/glands were mixed with Matrigel and seeded into 24-well tissue culture plate wells as described above.

### Total RNA extraction

Tissue: During post-mortem examination, tissue sections (0.5 x 0.5 cm) were excised from glandular tissue regions and placed into RNA*later™* solution (Fisher Scientific, 10391085). Tissue sections collected into RNA*later* were derived from tissue regions directly parallel to where tissue sections had been collected for organoid-generating material. Tissue samples in RNA*later* were stored at −70 °C until RNA extraction.

After defrosting at 5 °C, tissue was removed from RNA*later* solution onto a sterile petri dish using sterile forceps and a small piece (<10 mg) of the epithelial layer was carefully removed using a sterile scalpel and placed into prechilled RLT buffer containing β-mercaptoethanol (10 µl/ml, Thermofisher Scientific, Cat 125472500) in a Precellys® tissue homogenization tube containing 2.8 mm ceramic zirconium oxide beads (Bertin Technologies, P000911-LYSK0-A; CK28). Samples in Precellys tubes were then placed into a Bertin Precellys 24 tissue homogenizer and homogenized for 20 seconds at 5000 rpm. After homogenizing, samples were briefly centrifuged at 200 x *g* for 1 minute at 21 °C to remove bubbles and separate remaining tissue pieces to the bottom of the sample tube. Supernatant containing RNA was removed to a 1.5 ml tube and briefly mixed 1:1 with 70% ethanol diluted with RNAse-free water, then placed into a Qiagen RNEasy spin column.

Organoids: RNA was extracted from GL-organoids at passages P2 and P6, and organoids differentiated in OGM and ODM at P7. Organoids were gently removed from Matrigel® domes using ice-cold DMEM F/12, centrifuged at 300 x *g* at room temperature to pellet and resuspended in RNA*Later* solution. Samples were then immediately stored at −70°C until RNA extraction.

Total RNA was extracted from both tissue and organoids using a Qiagen RNeasy kit (Qiagen, ID.74104). Organoids were initially centrifuged at 500 x *g*, and RNA*Later* solution removed. Samples were then resuspended in RLT-buffer containing β-mercaptoethanol (10 µl/ml). Sample solution was mixed with 1:1 volume of 70% RNA-free ethanol and moved to a Qiagen RNeasy centrifuge column. From this step, manufacturer’s instructions were followed to extract RNA from organoids and tissue, including an on-column *DNAse-I* digestion for 15 mins at room temperature (Qiagen, ID.79254) prior to washing the column. All samples were eluted into 30 µl of nuclease-free water. Eluted RNA was quantified using a NanoDrop One spectrophotometer, and RNA integrity analysed on a Bioanalyser 2100 instrument using an RNA 6000 Nano Kit (Agilent technologies, 5067-1511). All samples used had an RNA integrity (RIN) value >6 and were stored at −70 °C until sequencing.

### RNA-seq analysis

Total RNA extracted from each sample (15 µg) was used for RNA-seq analysis. All library synthesis and sequencing was performed by Novogene (Cambridge, United Kingdom). In brief, dual-indexed, strand-specific RNA-seq libraries were constructed from submitted total RNA samples. The barcoded individual libraries were pooled and sequenced on an Illumina NovaSeq X Plus platform (Paired-end, 2×150 bp sequencing, polyA enrichment), generating on average 30 million reads per sample.

Upon retrieval of the data, reads were checked for quality using FastQC software (version 0.12) then pseudo-aligned to the *Equus caballus* genome (EquCab 3.0.113; retrieved from Ensembl database) using kallisto v.0.51.1 with default settings (Bray et al., 2016) with an average mapping rate of 77.6% matched reads. Reads were imported to RStudio (Posit Software & PBC, 2025; R Core Team, 2021), and genes with counts below 10 across all samples were removed to reduce the impact on the analysis of differentially expressed genes (DEGs). Reads were then analysed using DESeq2 package v.4.5 (Love et al., 2014) in R Studio to measure differential expression between tissue and organoids, organoids at P2 compared to P6 and organoids grown in OGM compared to ODM. Data was filtered using a Log_2_ Fold Change cutoff of 0.6 (corresponding to an approximately 1.5-fold change) and a false discovery rate (FDR)-adjusted p-value cutoff of <0.05 (Benjamini & Hochberg, 1995). Total-per-million (TPM) counts were log_2_ transformed after normalisation by DESeq2 and used to generate heatmaps using the“pheatmap” package. Biplots were generated using the “PCAtools” package (Blighe K & Lun A, 2025). Genes were annotated using the Ensembl database, using the “BiomaRt” package (Durinck et al., 2005, 2009), and the dataset “Equicab 3.0” (Kalbfleisch et al., 2018). Specific genes-of-interest were identified using the human protein atlas and published information (Supplemental Table 2) to identify cell-specific genes (Uhlén et al., 2015).

Glandular gastric tissue RNA-seq data for horse 2 was removed from DESeq2 analysis, as it was found to be an incorrect tissue (antral glandular gastric instead of corpus) and significantly impacted gene expression analysis. This discrepancy was initially identified by PCA-assay showing this tissue as an outlier, and further confirmed by absence of genes associated with parietal cells (*ATP synthase subunits 4a and 4b* (*ATP4A, ATP4B), aquaporin 4 (AQP4)* and *Cobalamin binding intrinstic factor (CBLIF*), chief cells (*PGA* isoforms) which are expressed in the gastric corpus but not in the antrum, and an increase in G-cell gene for *ghrelin* (*GHRL*) which are primarily found in the antrum. Organoids from this animal were left in analysis as they remained similar to all other tissue-derived organoids in gene expression.

Data visualisations were created using RStudio packages “ggplot2” (Wickham, 2016), and colour-blindness-friendly graphs created using the “viridis (Lite)” package v.0.6.5 (Garnier et al., 2024).

### Immunohistochemistry / Immunofluorescence

Tissue collected at post-mortem was immediately placed in 10% neutral buffered formalin (NBF) and stored for at least 24 hours until sufficiently fixed, then stored in 70% ethanol before being processed to wax and embedded in paraffin.

Gastric organoids were processed to a pellet, embedded in Histogel® (Epredia, HG-4000-012), processed to wax, and embedded in paraffin. Briefly, organoids were cultivated in Matrigel® and fed with OGM for 7 days. Once grown, feed media was removed and 1 ml of ice-cold Advanced DMEM/F12 was added to dissolve Matrigel® matrices. The suspended organoids were then transferred to a 15 ml Falcon tube and centrifuged at 200 x *g* for 5 minutes at room temperature. The supernatant was discarded, and organoids re-suspended in ice-cold 10% NBF. Organoids were incubated for 30 minutes at room temperature, then centrifuged at 400 x *g* for 5 minutes to pellet. The 10% NBF was removed and replaced with 10 ml PBS, briefly vortexed, then centrifuged at 300 *x g* for 5 minutes. PBS was removed until only a pellet remained, which was re-suspended in approximately 200 μl pre-warmed Histogel®, then added to a 7 x 7 x 5mm cryomold (Agar Scientific, AG27147-1) until set (approximately 10 minutes on ice). The Histogel® block was removed from the cryomold and placed into a cassette, then stored in PBS until processing to wax and paraffin embedding.

After paraffin embedding, both organoids and tissue blocks were sectioned to 5 µm thick sections using a Leica Jung Biocut 2035 microtome, adhered to Epredia™ Superfrost+™ slides, then dried overnight and stored at RT until use.

After sectioning, slides were deparaffinized and rehydrated using the following process. Slides were placed into 100% xylene for 3 minutes, fresh 100% xylene for 3 minutes, 2 dips into 1:1 xylene:ethanol, 100% ethanol for 3 minutes, 95% ethanol for 3 minutes, 70% ethanol for 3 minutes, then placed into tap water for 4 minutes. Antigen retrieval was then performed. Slides were completely submerged in sodium citrate (pH 6) buffer then heated in an autoclave to 90 °C for 15 minutes. After being allowed to cool at room temperature, slides were rinsed 3 times in Tris-buffered saline (TBS). Tissue was permeabilized using 0.1% triton X-100 in TBS for 10 minutes at room temperature, then blocked for 1 hour at room temperature using 10% BSA in TBST (TBS-0.25% Tween 20). Primary antibodies dissolved in blocking buffer were added to organoid slides and incubated overnight at 4 °C. The antibodies were used at the following dilutions; Rabbit anti-POU2F3 (Sigma Aldrich, HPA01985; 1/200) to identify tuft cells, rabbit anti-Ki67 (Abcam ab15580; 1/1000) to identify replicating cells, mouse anti-ZO-1 (Fisher Scientific, 10129012, 1/500) to identify tight-gap junctions, rabbit anti-Mucin 5ac (Abcam, ab3649, 1/200) to identify foveolar cells. An antibody against chromogranin A (ChgA) was used to identify EECs (rabbit anti-chromogranin A (NovusBiologicals, NB120-15160; 1/500). Slides were then washed 3 times using TBST. Secondary antibody (Goat anti-rabbit IgG AlexaFluor™ 488 (Invitrogen, A-11008) or Goat anti-mouse IgG AlexaFluor™ 555 (Invitrogen, A-21422)) was then diluted in blocking buffer and placed on slides for 1 hour at room temperature in the dark. Slides were subsequently washed 3x with TBST. Finally, DAPI (1 µg/ml; Invitrogen, D1306) was applied to slides for 10 minutes at RT in the dark. Slides were then mounted using ProLong Gold antifade medium (Invitrogen, P36934) and allowed to set overnight at 5 °C in darkness. Slides were then sealed using clear commercially available nail varnish.

Periodic-acid-schiff (PAS) staining was performed by deparaffinising tissue as previously described, then oxidising slides using 1% periodic acid for 5 minutes. Sections were rinsed in tap water for 5 minutes, then treated with Schiff’s reagent (TCS bioscience; HS265-500) for 15 minutes. Slides were rinsed under running tap water for 20 minutes followed by being submerged in haematoxylin-Z (Cellpath; RBA-4201-00A) counterstain for 3 minutes, before again rinsing in tap water until clear. Slides were briefly dipped in 1% acid-alcohol to remove excess haematoxylin, then nuclei blued using Scott’s tap water substitute for 30 seconds. Finally, slides were dehydrated using the reverse procedure for deparaffinisation, then coverslips mounted using Epredia ClearVue mountant (Fisher Scientific; 11396790).

### Imaging and data analysis

Brightfield and DIC images of organoids were taken on a Zeiss Axiovert 200 M microscope using an AxioCamMR3, and a Colibri 7 LED light source, operating with Zen Blue software (version 3.1) Fluorescent images of organoids were taken on a Zeiss AxioObserver using a plan-Apochromat 20x objective and Axiocam 705 camera.

## Results

### Establishment of equine gastric organoids

Gastric glands were successfully isolated from fundic mucosal tissue (Figure 3.A). After seeding in Matrigel®, glands were observed daily for the development of self-organizing GL-organoids. These organoids were observed to develop within 24 hours and rapidly formed a branching phenotype with a thin lumen (Figure 3.A, P0), similar to the glandular starting material. No significant changes in phenotype were observed as organoids were passaged to P2.

**Figure 3:**
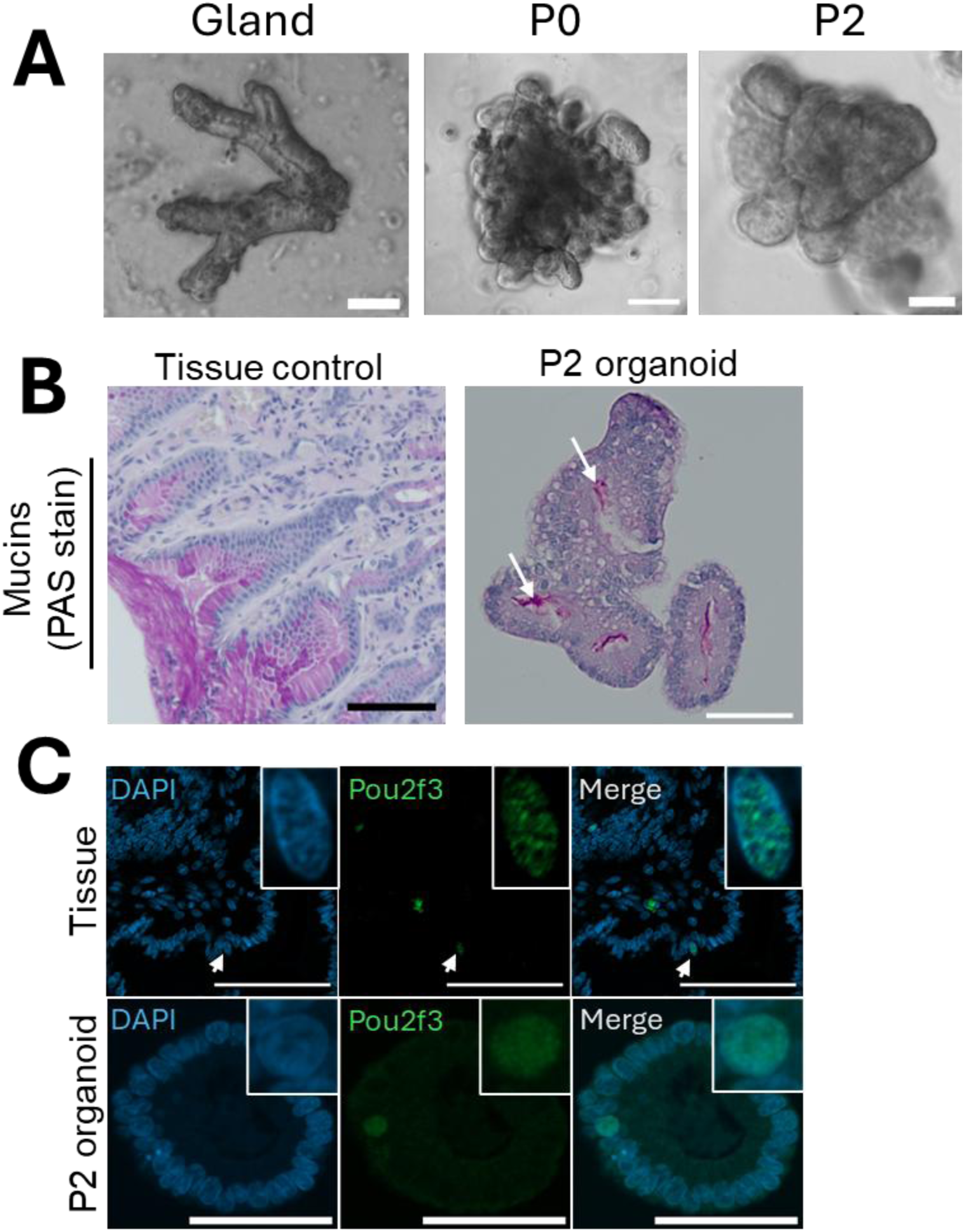
Establishing GL-organoids from tissue glands. **A:** gastric glands digested from tissue showed expected branching morphology. At P0 and P2, GL-organoids derived from these glands retained a branching phenotype with a clear lumen. **B:** PAS stains of equine glandular gastric tissue (pit region) showing strong mucus production (left, bright magenta coloured areas) and P2 GL-organoid showing mucus released into lumen. **C**: Immunofluorescence showing rare Pou2f3+ tuft cells in glandular gastric tissue (top) and P2 organoids (bottom). All organoids collected and processed for imaging after 5 days of growth. Scale bars represent 50 µm.

A well-established characteristic of gastrointestinal organoids in other species is the presence of multiple epithelial cell types. PAS staining of equine glandular gastric organoid sections revealed the presence of a mucus layer within the lumen of the organoids, a characteristic shared with the apical surface of the gastric mucosa *in vivo* and indicating the presence of mucus-producing cells (Figure 3B). However, another secretory lineage cell type was found to be extremely rare in organoids cultured in expansion conditions, with only a single cell nucleus labelling positively for the tuft cell-specific marker Pou2f3 (Figure 3C), despite testing multiple organoid sections derived from each animal in the study, each of which contained multiple organoids.

Having successfully established that GL-organoids can be derived from equine gastric tissue, we next sought to determine the long-term stability of these organoids in culture by passaging an additional 4 times to P6. We found that at P6 the GL-organoids retained their branching, luminal phenotype (Figure 4A) and continued to secrete mucus into the lumen (Figure 4B).

**Figure 4:**
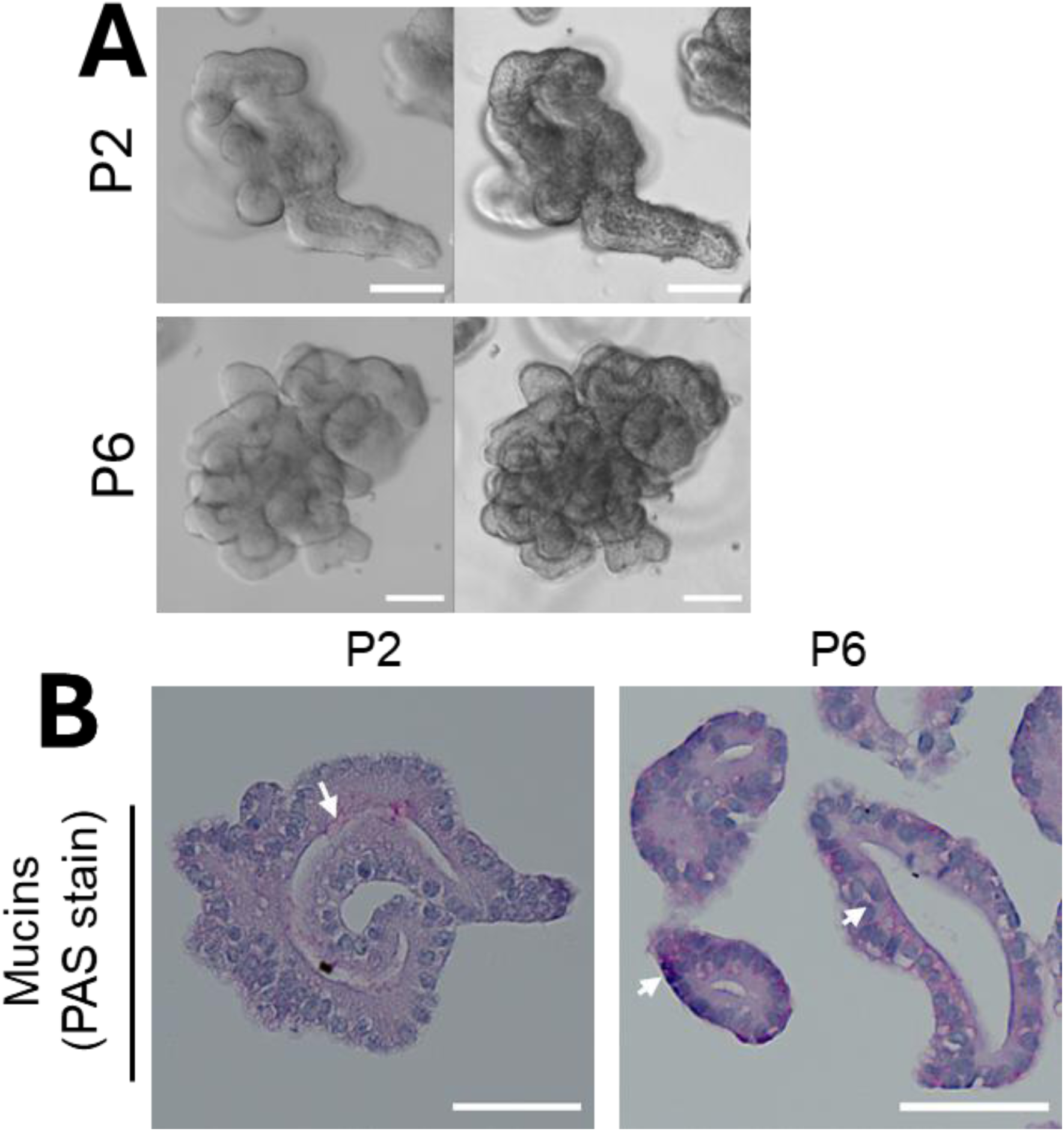
Equine GL-organoids can be passaged to P6. A: GL-organoids retain a branching, luminal phenotype when passaged from P2 (left) to P6 (right). B: GL-organoids produce mucus at P2 and P6. All GL-organoids were imaged at day 5, all scale bars represent 50 µm.

Next, we wanted to determine if the GL-organoids could be differentiated to express differentiated, specialized epithelial cell-types. After differentiation for 4 days in ODM media, organoids from each animal formed a swollen appearance with a wide lumen and loss of the branching phenotype (Figure 5A). ODM-cultured organoids retained tight junction (ZO-1) labelling across all cell-to-cell boundaries and organoids were still found to contain few proliferative (Ki67-positive) cells (Figure 5B). This phenotype was observed to change gradually over the 4 days and was associated with an increased differentiation of mucus-producing cells, with PAS staining revealing an increase in both intracellular mucus, as well as mucus secreted within the organoid lumen (Figure 6C). Subsequent IF labelling confirmed the tissue-specificity of the mucus produced in the GL-organoids, with only differentiated organoids labelling positively for the gastric-specific mucin Mu5ac (Figure 6D).

**Figure 5:**
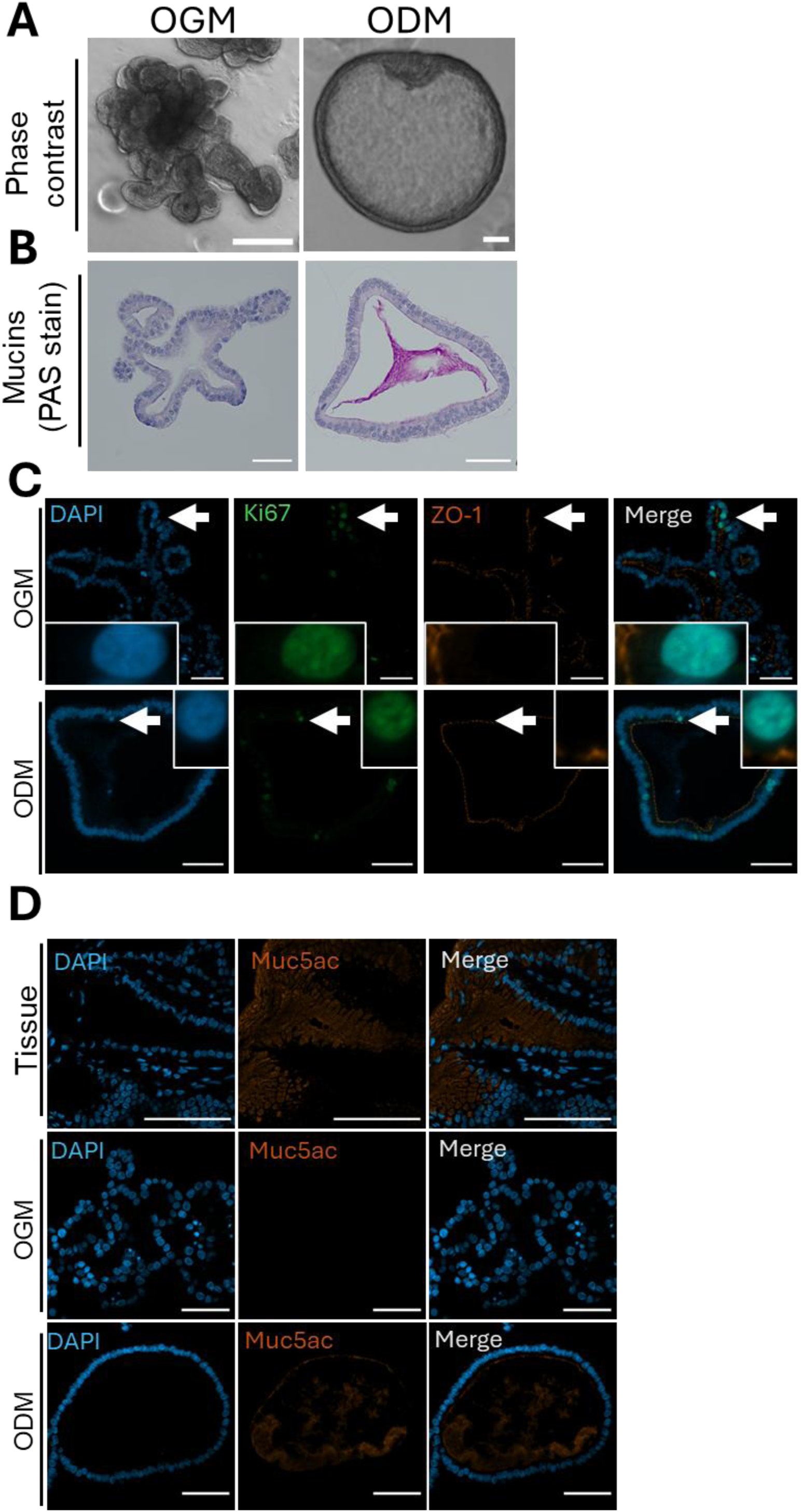
Differentiation of equine GL-organoids. Equine GL-organoids were grown in differentiation media for 4-5 days before being collected for IF/PAS stains. **A**: Phase contrast images of GL-Organoids grown in OGM (left) and ODM (right). Change in phenotype occurred gradually over 4 days. **B**: PAS stain showing mucus production of GL-organoids grown in OGM and ODM. **C**: IF stain for replicating cell marker Mki67 and tight-gap junction marker ZO-1 in organoids grown in OGM and ODM. **D**: IF stain for gastric-specific mucus Muc5ac shown in the pit region of the gastric gland in tissue, organoids grown in OGM and organoids grown in ODM. All scale bars represent 50 µm.

**Figure 6:**
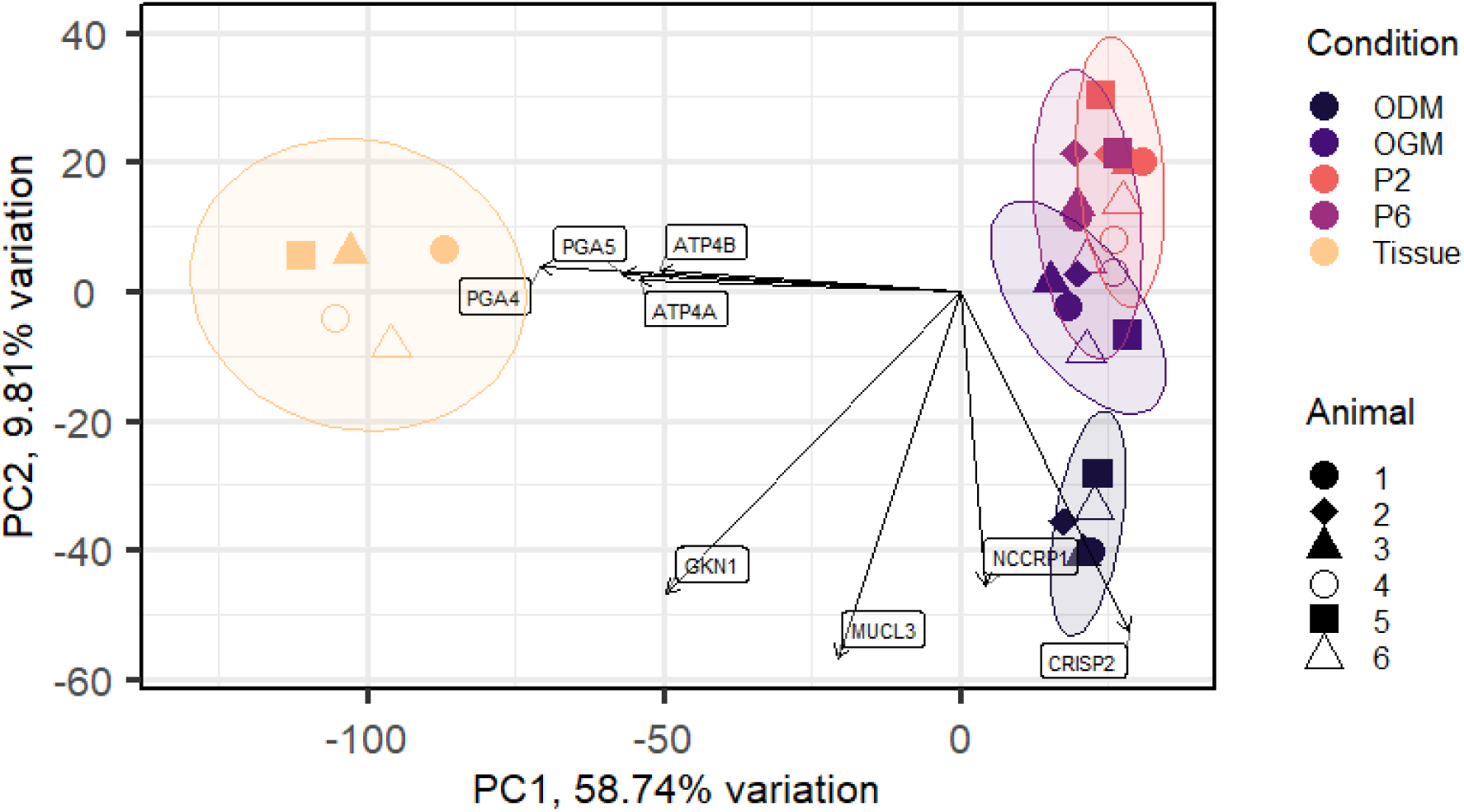
Biplot showing principal component analysis of the top 500 most variant genes. Genes are ranked by inter-sample variance in equine glandular gastric tissue and organoids, and the key genes influencing these components. Different colours show different conditions, and different shapes show different animals. Ellipses show groupings at 95% confidence interval.

In order to provide a comprehensive analysis of the equine GL-organoids, bulk RNA-sequencing analysis was performed on organoids and the tissue they were derived from. The global gene expression profile of GL-organoids compared to tissue was first analyzed by principal component analysis (PCA) and a biplot generated to show the key genes influencing these variations (Figure 7). The PCA analysis showed grouping of tissue primarily influenced by expression of parietal and chief cell genes *PGA*, *ATP4A* and *ATPB*, with P2, P6 and OGM GL-organoids grouping together and the ODM condition forming a third distinct group influenced by expression of surface mucus cell-associated mucin-like 3 (*MUCL3), gastrokine 1 (GKN1)*, cysteine-rich secretory protein 2 (*CRISP2)* and F-box associated domain containing protein (*NCCRP1)*. Both tissue and organoid groupings showed relatively similar gene expression between individual animals despite the wide variation in breed, age and sex.

**Figure 7:**
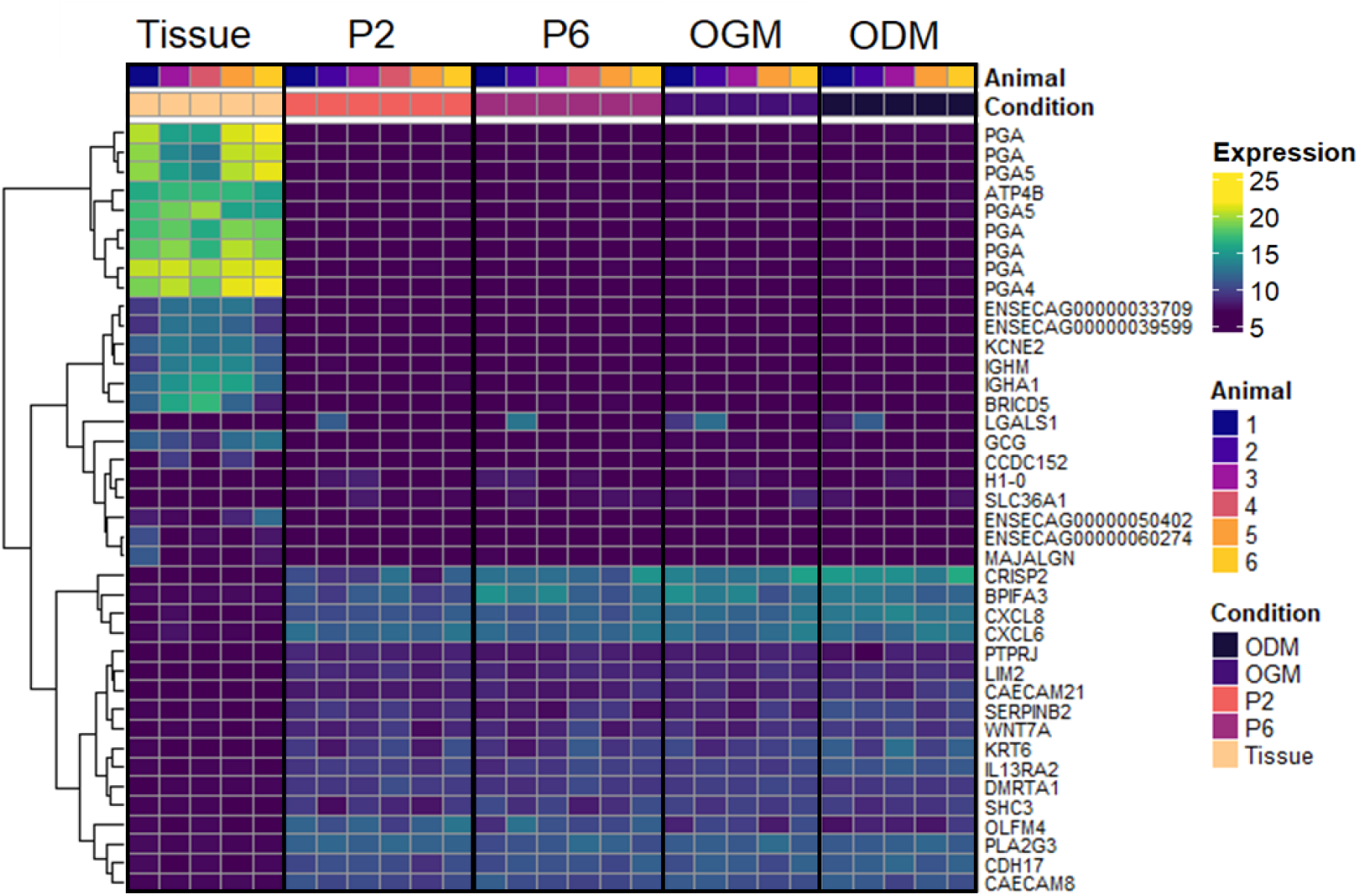
Top 40 differentially expressed genes of glandular gastric tissue and organoids. Heat map showing expression level of the top 40 most differentially expressed genes (ranked by inter-sample variance) from equine glandular gastric tissue and organoids derived from that tissue (n=5) at p2 (n=6), P6 (n=6) and grown in OGM (n=5) and ODM. (n=5). Colours indicate level of expression low (dark, purple) to high (light, yellow). Dendograms indicate similarity between sample groups. Gene symbols annotated from ensemble database where possible. Details of genes included in the heatmap are shown in Supplemental Table 1. Colourblindness-friendly colour scale “viridis” used.

The expression profile of the top 40 most variable genes between GL-gastric tissue and GL-organoids (all conditions) were then compared. This analysis showed distinct grouping of genes upregulated in tissue as compared to organoids, and genes downregulated in tissue as compared to organoids (Figure 8). It was found that the majority of upregulated DEGS between tissue and organoids were *PGA* isoforms which encode for pepsinogen produced primarily by chief cells in the gastric gland, as indicated by the biplot in figure 6. Transcripts upregulated in organoids varied, with the strongest differences in gene expression occurring in *CRISP2* (Cysteine-rich secretory protein 2), *BPIFA3* (BPI fold containing family A, member 3) and the chemokines *CXCL6* and *CXCL8* (Chemokine (C-X-C motif) ligands 6 and 8).

**Figure 8.**
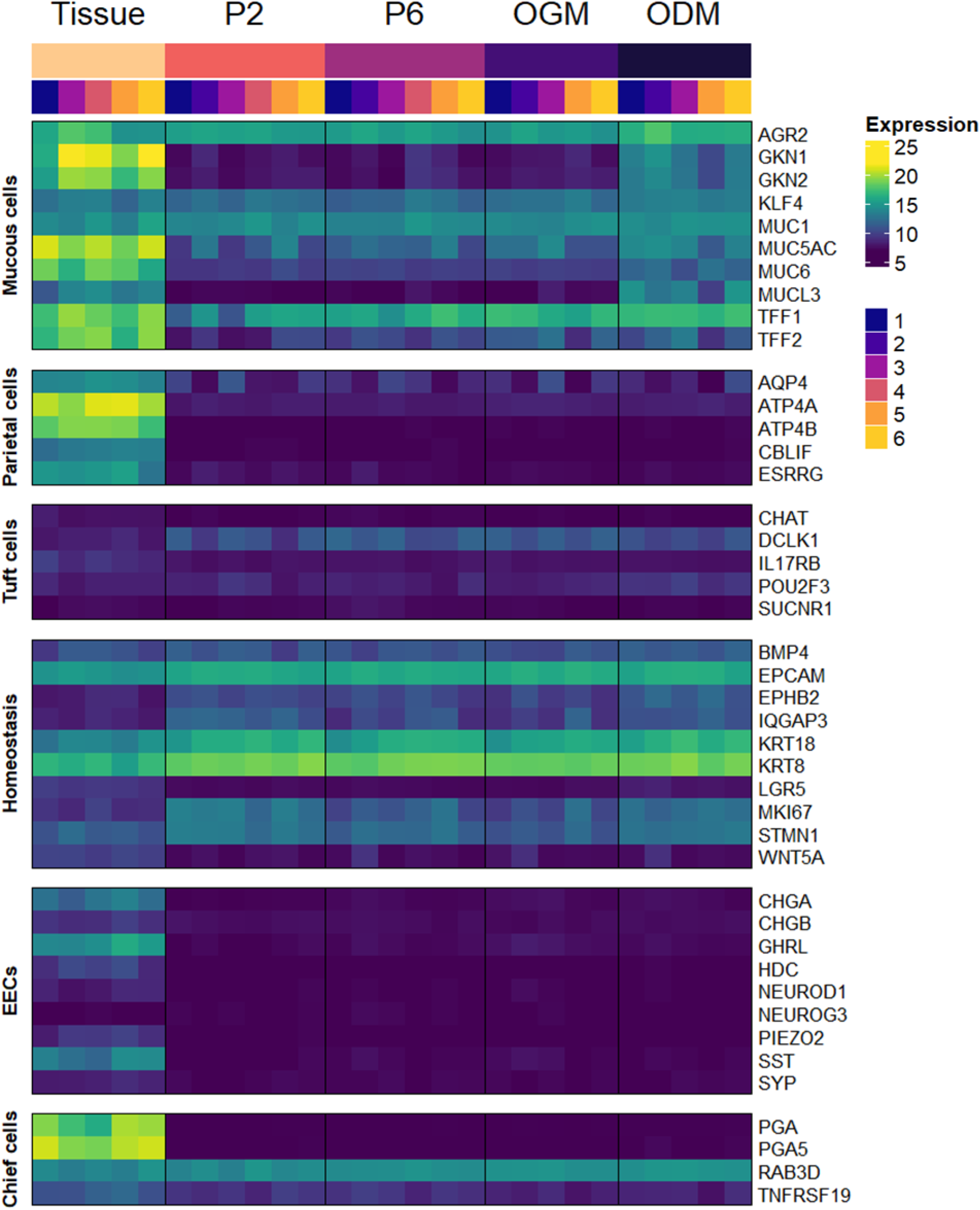
Cell specific gene expression of glandular gastric tissue and organoids. Heat map showing expression level of genes associated with differentiated cells of the gastric epithelium from equine glandular gastric tissue and organoids derived from that tissue (n=5) at P2 (n=6), P6(n=6) and grown in OGM (n=5) and ODM. (n=5). Colours indicate level of expression low (dark, purple) to high (light, yellow). Genes associated with specific cell-types and tissue-types identified using the Human Proteome Atlas database. Gene symbols annotated from ensemble database where possible. Details of genes included in the heatmap are shown in Supplementary Table 1.

Gene expression profiles from GL-organoids at P2 (n=6), P6 (n=6), OGM (n=5) and ODM (n=5) were compared to GL-gastric tissue (n=5) by RNA-seq analysis.

After filtering for low-expression genes (<10 counts), 8,105 significantly differentially expressed genes (DEGs) were found between P2 organoids and glandular gastric tissue. A similarly high number of DEGs (7,553) were identified between P6 organoids and tissue. In each comparison many genes were associated with non-epithelial cells, such as mu heavy constant chain of IgM(*IGHM)*, alpha-1 heavy constant chain of IgA (*IGHA1)* and epsilon heavy-constant chain of IgE (*IGHE),* however many of the most highly expressed genes that were downregulated in organoids were associated with chief and parietal cells, such as pepsinogen (*PGA*) isoforms, *ATP4A/B*, ghrelin (*GHRL*) and gastrokines (*GKN1*/2). Genes upregulated in organoids compared to tissue were keratins (*KRT6*, *KRT8*), lysozyme (*LYZ*), galectin 1 (*LGALS*1)-although this was mostly influenced by the organoid sample derived from glandular corpus tissue.

A transcriptomic analysis of organoid stability over time revealed 353 genes to be differentially expressed between P2 and P6 organoids cultured in OGM expansion media. Genes downregulated between P2 and P6 included Collagen type III alpha 1 chain (COL3A1) and genes associated with nutrient uptake such as solute carrier family genes *SLC7A5, SLC38A5* and *SLC3A2.* Upregulated genes included *SLC37A1*, *SLC16A12* and *SLC29A4* and EEC gene *CHGA*. Of note, the most differentially upregulated gene in P6 organoids compared with P2 was the equine allergen protein C1 (*EQUC1*) with a log2FC of 15.2.

A pairwise comparison of gene expression between organoids cultured in OGM and ODM determined differential expression in 1917 genes. Key genes upregulated in ODM were associated with the gastric epithelium such as gastrokines (*GKN1*, *GKN2*), mucins (*MUCL3*, *MUC6*) and replication marker *MKI67*. Downregulated genes included polymeric immunoglobin receptor (PIGR) and several eukaryotic elongation factors (*EEF1A1, EEF2, EEF1G*). Full lists of DEG for each comparison can be found in supplemental data 1.

**Table 2:** Summary of upregulated and downregulated differentially expressed genes in each DeSeq2 comparison.

| Comparison | Total DEGs |
| --- | --- |
| P2 organoids vs Tissue | 8105 |
| P6 organoids vs Tissue | 7533 |
| P2 vs P6 Organoids | 353 |
| OGM vs ODM organoids | 1917 |

Next, we sought to use the gene expression data to determine which gastric gland cell types were likely to be present within the GL-organoids, across passages and in growth vs differentiation conditions. Initially, cell type-specific gene transcript markers were identified from the published literature (Supplementary File 1). An analysis of the transcript counts for these markers showed that while mucous genes were consistently expressed in both tissues and organoids, many were downregulated in P2 and P6 organoids as compared to tissue (*MUC5AC, MUC6*, *TFF1, GKN1, GKN2*), consistent with Muc5ac antibody labelling in organoids grown in OGM (Figure 5D). However, ODM-cultured GL-organoids increased expression of these genes, indicating differentiation towards gastric pit cells (Figure 8) While tuft cells were difficult to detect by antibody labelling (Figure 3), the expression of the key tuft cell transcript markers *DCLK1* and *POU2F3* was identified in organoids from all animals and under all conditions. GL-organoids did not strongly express any genes associated with EECs, consistent with attempts to antibody label for the EEC marker ChgA which was consistently found to be negative in all organoid sections (data not shown). We also did not find expression of *ATP4A/B,* characteristic of parietal cells, nor any isoforms of the *PGA* genes of chief cells, suggesting the conditions were not optimal for expression of these specialized epithelial cells. Many genes associated with homeostasis such as *EPCAM*, *EPHB2*, *BMP4* were consistent between tissue and organoids, or even increased in organoids such as *MKI67 and STMN1* suggesting that even under differentiation conditions the organoids continued to express replicating-cell markers, consistent with Ki67 antibody labelling (Figure 5).

## Discussion

One of the biggest problems faced by equine research is the difficulty and expense of using live animals in research. Horses are large, expensive to keep, and in the UK are classified as companion animals making research on horses challenging and ethically questionable. Given how many horses worldwide are estimated to suffer from a variety of gastric diseases (Alloway et al., 2020; Sequeira et al., 2001; Vokes et al., 2023) there is a need to develop a means of performing *in vitro* research, which can accurately recapitulate the diversity of signaling and cell-types of the equine gastric epithelium.

This study describes the establishment of a first-of-its-kind organoid model of the equine GL-gastric epithelium that will expand the capabilities of equine researchers to understand diseases and infections of the equine stomach. Previous research has been conducted using live animals, and to our knowledge there is no equine gastric epithelial cell line commercially available. We have shown that by isolating the stem cell-rich glands from equine gastric tissue, multi-cellular organoids can be cultured long-term and that these organoids retain their gene expression profile and phenotype over multiple passages, consistent with gastric organoid cultivation for other mammalian species. Also consistent with other gastrointestinal organoid models and important to the sustained cultivation of this epithelial model was the presence of proliferative markers (e.g. Ki67) and tight junction markers (e.g. ZO-1). Of significant importance is the finding that our organoids showed little animal-to-animal variation in transcriptomic responses despite the variation in the age, breed, sex and clinical presentation of the horses used for this study, indicating that horses from diverse backgrounds can be used for organoid generation with minimal anticipated variation in organoid phenotype.

While organoid models have been developed from the equine intestinal tract, such as the jejunum (Hellman et al., 2024; Stewart et al., 2018) and colon (Windhaber et al., 2024), to our knowledge this is the first time organoids have been derived from the equine glandular gastric mucosa. Similarly, this paper describes the first transcriptomic analysis of equine organoids of any type and is the first to use transcriptomics to comprehensively show these organoids remain representative of the source tissue over multiple passages, as well as determine the key differences between sample types described here.

A major component of equine organoids derived from gastric glands was mucous-producing cells, as evidenced by mucin gene expression and the presence of mucus. Furthermore, this mucus expression was enhanced in organoids cultured in a differentiation media. The mucin types expressed in the glandular organoids confirmed their glandular phenotype specificity (Bell et al., 2007; Bullimore et al., 2001). More specifically, our data suggests that application of Intesticult™ ODM pushes organoids to a gastric pit-type phenotype, with increased expression of genes such as *MUC5AC* and *GKN1/2* (associated with gastric foveolar cells; (Babu et al., 2006) and *MUC6* and *MUCL3* (associated with mucous neck cells; (Kang et al., 2005)). While transcripts were present for *MUC5AC* in P2, P6 and OGM organoids, the corresponding protein was not detectable by IF without cultivation in ODM. The differentiation of equine gastric organoids towards a gastric pit cell type after the withdrawal of Wnt also suggests the signaling in the horse glandular gastric epithelium may be similar to other species (Hong et al., 2025).

Based on a combination of histological and transcriptomic analysis, other differentiated cell types such as tuft cells were identified within equine glandular gastric organoids, albeit at much lower abundance. A notable difference observed between our GL-organoids and biopsied gastric tissue was the absence of chief cells, parietal cells and EECs. This is potentially explained by the fact that our organoids consistently expressed *MUC5AC* and *TFF1* suggesting they were exhibiting a gastric pit genotype even before differentiation, and EECs, parietal cells and chief cells are not found in the gastric pit, but instead are found deeper within the gastric gland in the isthmus and base (Wölffling et al., 2021). The absence of EECs and low numbers of tuft cells in equine gastric organoids after differentiation is in contrast to those previously published using organoids derived from the equine jejunum and colon (Windhaber et al., 2024). This is likely due to the key differences in cell signaling between the gastric and small intestine mucosa, and emphasizes the importance of adapting media conditions to the source tissue and desired outcome. Methods of adapting media conditions to differentiate to these cell types within gastric organoids have previously been published for humans and mice (Adkins-Threats et al., 2024; Hong et al., 2025; Wölffling et al., 2021), which can be used as a baseline for further adaptation of this model to facilitate research into specific cells and regions of the equine stomach.

It is important to note that a major difference between the transcriptome of *ex vivo* tissue and organoids is expression of genes not associated with the epithelium (e.g. genes associated with the immune system, and submucosa). This is to be expected, as mammalian gastrointestinal organoids are typically a model of the surface epithelium alone and do not express multiple tissue-types (e.g. musculature, submucosal immune cells or neurons), while *ex vivo* tissue is a highly complex mix of different structures and tissue types, and *in vitro* these cell types either do not survive the initial digestion or do not remain in culture after multiple passages due to selective media conditions (Faber et al., 2022; Smith et al., 2021).

## Conclusions

Overall, this study describes the methods used to establish, characterize and differentiate equine glandular gastric organoids. We have demonstrated that this *in vitro* model is physiologically relevant and that gene expression and phenotype are retained across multiple passages, meaning that these organoids can be cultured for extensive periods of time without undergoing significant phenotypic changes. We show these organoids can be differentiated to better represent specific cell-types within the gastric mucosa. This makes equine gastric organoids a highly adaptive, stable and applicable tool for researching equine gastric diseases and therapies, reducing dependency on live animal studies to investigate biology of the equine gastric epithelium.

## Declarations

### Ethical approval

This study was performed under MRI Equine Grass Sickness Biobank AWERB approval 160321. Veterinary surgeons involved in euthanasia obtained owner/client permission to post-mortem and extract tissue to be used in research. Veterinary surgeons signed declaration forms that owner/client permission has been obtained.

### Data availability

Data supporting the findings described in this manuscript are available in the supplementary materials.

### Author contributions & competing interests

Tanith Harte developed methodology, generated and analyzed data, and drafted this manuscript. Tanith Harte, David Smith and Beth Wells conceptualized and acquired funding. Helen Todd contributed to immunofluorescence data generation. Beth Wells, Jo Moore, Stewart T.G. Burgess and David Smith edited and critically reviewed this manuscript. All authors read and approved the final manuscript.

The authors declare that they have no competing interests.

