## Supplementary figures and images for "Development and Characterisation of Equine Gastric Organoids"

### Supplemental figure 1

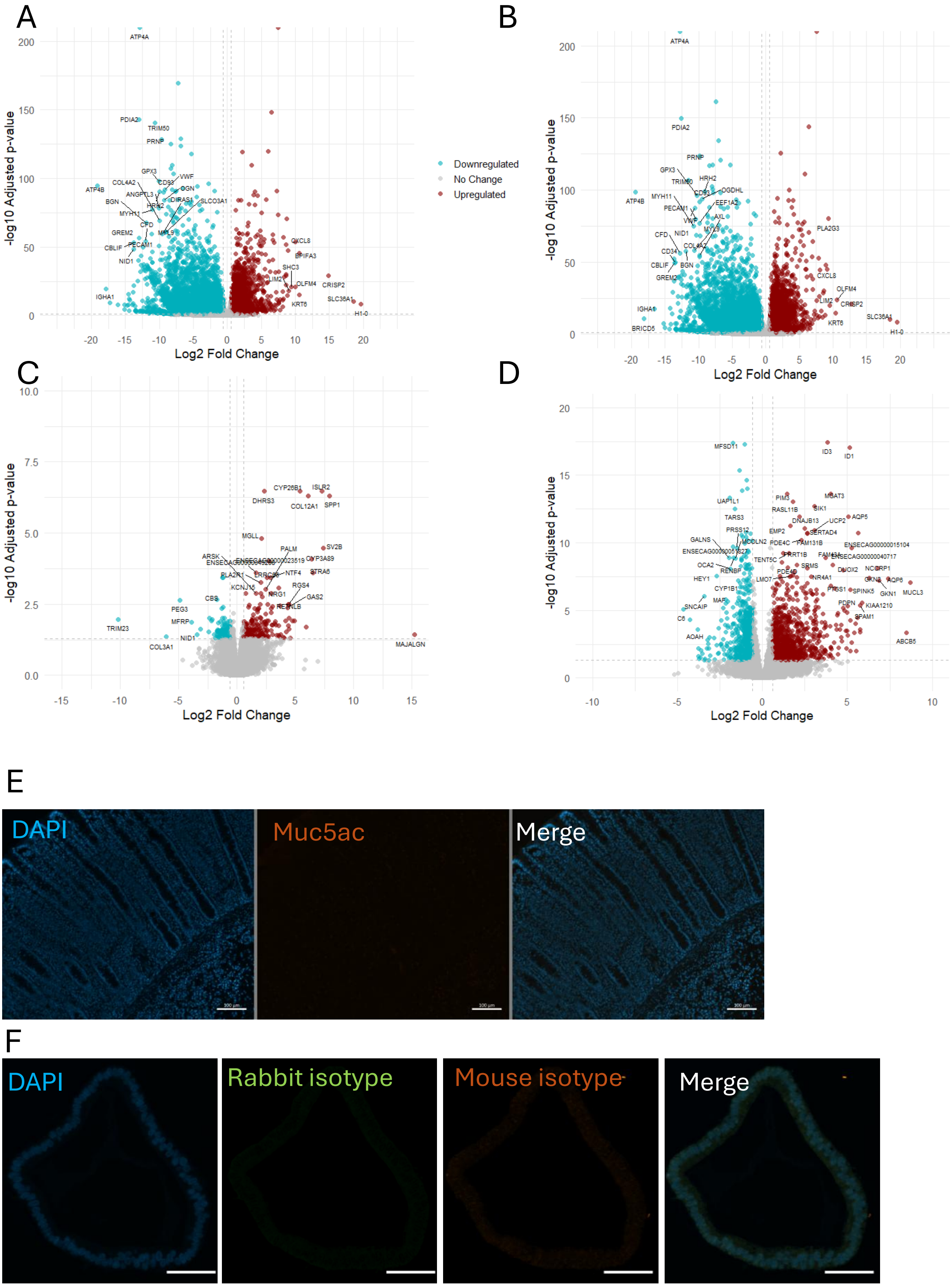
