## Supplemental table 1 for "Development and Characterisation of Equine Gastric Organoids"

Supplementary tables

Table 1: Materials, reagents and supplier information.

| **Supplier** | **Product Name** | **Product Code** |
| --- | --- | --- |
| **Organoid Culture Reagents** | | |
| Stemcell Technologies^TM^ | Intesticult^TM^ Organoid Growth Media (Mouse) | 0600 |
|  | Intesticult^TM^ Organoid differentiation media (Human) | 100-0214 |
|  | Cryostor CS10 | 100-1061 |
| Fisher Scientific | Gibco HBSS with calcium and magnesium | 11560616 |
|  | Gibco N2 supplement (50x) | 11520536 |
|  | Gibco B-27 supplement (100x) | 12587010 |
|  | Gibco Advanced DMEM F/12 | 11540446 |
|  | Gibco DMEM High glucose, pyruvate | 13345364 |
|  | Invitrogen RNA*later* stabilization solution | 10391085 |
|  | Corning® Matrigel®GFR Basement membrane matrix | 11543620 |
|  | Cytiva HyClone Amphotericin B Solution | 11526481 |
|  | Gibco Gentamicin (50mg/ml) | 11530506 |
|  | Bertin Technologies Hard tissue homogenizing 2ml kit (CK28) | 11598555 |
| MRI stores | Pencillin / Streptomycin solution |  |
|  | Trypsin |  |
|  | Versene |  |
| InvivoGen | Normocin® | Ant nr-05 |
| Cambridge Bioscience | SB-202190 | CAY10010399 |
|  | LY2157299 | CAY15312-10mg |
| Tocris Biotechne | Y-27632 Dihydrochloride | 1254 |
| Roche | Dispase | 4942086001 |
| Merck/Sigma Aldrich | Collagenase | C2674 |
| **Histology / Immunofluorescence reagents** | | |
| Fisher Scientific | Epredia HistoGel Specimen Processing Gel | 12006679 |
|  | Invitrogen ProLong Gold Antifade Mountant | 11539306 |
|  | Invitrogen DAPI Nucleic Acid Stain | 10184322 |
|  | Epredia ClearVue Mountant | 11396790 |
| MRI Stores | Phosphate buffered saline (PBS) |  |
| Cellpath | Hematoxylin-Z | RBA-4201-00A |
| VWR Chemicals | Xylene | 29875.360 |
| TCS Bioscience | Schiff’s reagent | HS265-500 |
| Agar Scientific | Disposable Base Moulds 7x7x5mm | AG27147-1 |
| **Antibodies** | | |
| Merck/Sigma Aldrich | Rabbit anti-POU2F3 | HPA019652-100UL |
| NOVUS Biologicals | Rabbit anti-Chromogranin A | NB120-15160SS |
| Invitrogen | Mouse anti-ZO-1 | 33-9100 |
| Abcam | Rabbit anti-Ki67 | Ab15580 |
| Abcam | Mouse anti-Muc5ac | Ab3649 |
| Invitrogen | Goat anti-rabbit IgG AlexaFluor™ 488 | A-11008 |
|  | Goat anti-mouse IgG AlexaFluor™ 555 | A-21422 |
| **RNA extraction reagents** | | |
| Qiagen | RNEasy Mini Kit | 74104 |
|  | RNase-Free DNase Set | 79254 |
| Invitrogen | RNA*later*™ Stabilization Solution | AM720 |
| Thermofisher Scientific | 2-Mercaptoethanol, 99%, pure | 125472500 |
| Bertin Technologies | Precellys® Hard tissue homogenizing CK28 – 2 mL | P000911-LYSK0-A |
| Agilent technologies, | RNA 6000 Nano Kit | 5067-1511 |

Supplementary Table 2: Cell-specific genes used to identify presence of differentiated cell-types in equine GL-organoids

Cell-specific gene expression was used to identify the specific cell types within the tissue and organoid transcripts. Cell-specific genes were identified by reviewing previous published information on the cell-type in combination with the human protein atlas (proteinatlas.org).

| *Associated cell-type* | *Gene name* | *Gene ID* | Reference |
| --- | --- | --- | --- |
| Mucous (foveolar / mucous neck) | Gastrokine 1 & 2 | *GKN1 / GKN2* | *(Kim et al., 2014)* |
|  | Mucin 5ac | *MUC5AC* | *(Alvina et al., 2023)* |
|  | Mucin 6 | *MUC6* | *(Uhlén et al., 2015)* |
|  | Trefoil factor 1 & 2 | *TFF1 / TFF2* | *(Alvina et al., 2023)* |
| Mucous (other) | Mucin 1 | *MUC1* | *(Jin et al., 2017)* |
|  | Mucin-like 3 | *MUCL3* | *(Hong et al., 2025)* |
|  | Kruppel-like factor 4 | *KLF4* | *(Alvina et al., 2023)* |
|  | Anterior gradient protein 2 | *AGR2* | *(Gupta et al., 2013)* |
| Tuft cells | Choline acetyltransferase | *CHAT* | *(Hildersley et al., 2021)* |
|  | Doublecortin-like kinase 1, | *DCLK1* | *(Hildersley et al., 2021)* |
|  | Interleukin 17 receptor B | *IL17RB* | *(Feng et al., 2025)* |
|  | POU class 2 homeobox 3 | *POU2F3* | *(Hildersley et al., 2021)* |
|  | Succinate receptor 1 | *SUCNR1* | *(Hildersley et al., 2021)* |
| EECs | Chromogranin A & B | *CHGA / CHGB* | *(Busslinger et al., 2021)* |
|  | Ghrelin | *GHRL* | *(Alvina et al., 2023)* |
|  | Histamine decarboxylase | *HDC* | *(Latorre et al., 2016)* |
|  | Neurogenic differentiation 1 | *NEUROD1* | (Busslinger et al., 2021) |
|  | Neurogenin-3 | *NEUROG3* | *(Alvina et al., 2023)* |
|  | Piezo2 | *PIEZO2* | *(Mercado-Perez et al., 2024)* |
|  | Somatostatin | *SST* | *(Latorre et al., 2016)* |
|  | Synaptophysin | *SYP* | (Portela-Gomes et al., 1999) |
| Chief cells | Pepsinogen | *PGA (many isoforms)* | *(Alvina et al., 2023)* |
|  | Tumor necrosis factor receptor superfamily member 19 | *TNFRS19* | *(Stange et al., 2013)* |
|  | Ras-related protein 3D | *RAB3D* | *(Tang et al., 1996)* |
| Parietal cells | Aquaporin 4 | *AQP4* | *(Carmosino et al., 2005)* |
|  | Cobalamin binding intrinsic factor | *CBLIF* | *(Alvina et al., 2023)* |
|  | Estrogen-related receptor gamma | *ERRG* | *(Adkins-Threats et al., 2024)* |
|  | ATPase H+/K+ transporting subunit alpha & beta 4 | *ATP4A / ATP4B* | *(Alvina et al., 2023)* |
| Homeostasis | Bone morphogenic protein 4 | *BMP4* | *(Hong et al., 2025)* |
|  | Epithelial cell adhesion molecule | *EPCAM* | *(Matsuo et al., 2021)* |
|  | Ephrin type-B receptor 2 | *EPHB2* | (Perez White & Getsios, 2014) |
|  | IQ motif containing GTPase activating protein 3 | *IQGAP3* | *(Matsuo et al., 2021)* |
|  | Keratin 18 | *KRT18* | *(Kalabusheva et al., 2023)* |
|  | Keratin 8 | *KRT8* | *(Kalabusheva et al., 2023)* |
|  | Leucine-rich repeat-containing G-coupled protein receptor 5 | *LGR5* | *(Alvina et al., 2023)* |
|  | Ephrin type-B receptor 2 | *EPHB2* | (Perez White & Getsios, 2014) |
|  | Antigen Kiel 67 | *KI67 (MKI67)* | *(Han et al., 2019)* |
|  | Stathmin 1 | *STMN1* | *(Han et al., 2019)* |
|  | Wnt family member 5a | *WNT5A* | *(Alvina et al., 2023)* |

Adkins-Threats, M., Arimura, S., Huang, Y.-Z., Divenko, M., To, S., Mao, H., Zeng, Y., Hwang, J. Y., Burclaff, J. R., Jain, S., & Mills, J. C. (2024). Metabolic regulator ERRγ governs gastric stem cell differentiation into acid-secreting parietal cells. *Cell Stem Cell*, *31*(6), 886-903.e8. https://doi.org/10.1016/j.stem.2024.04.016

Alvina, F. B., Chen, T. C.-Y., Lim, H. Y. G., & Barker, N. (2023). Gastric epithelial stem cells in development, homeostasis and regeneration. *Development*, *150*(18). https://doi.org/10.1242/dev.201494

Busslinger, G. A., Weusten, B. L. A., Bogte, A., Begthel, H., Brosens, L. A. A., & Clevers, H. (2021). Human gastrointestinal epithelia of the esophagus, stomach, and duodenum resolved at single-cell resolution. *Cell Reports*, *34*(10), 108819. https://doi.org/10.1016/j.celrep.2021.108819

Carmosino, M., Mazzone, A., Laforenza, U., Gastaldi, G., Svelto, M., & Valenti, G. (2005). Altered expression of aquaporin 4 and H /K+‐ATPase in the stomachs of peptide YY (PYY) transgenic mice. *Biology of the Cell*, *97*(9), 735–742. https://doi.org/10.1042/BC20040138

Feng, X., Andersson, T., Flüchter, P., Gschwend, J., Berest, I., Muff, J. L., Lechner, A., Gondrand, A., Westermann, P., Brander, N., Carchidi, D., De Tenorio, J. C., Pan, T., Boehm, U., Klose, C. S. N., Artis, D., Messner, C. B., Leinders-Zufall, T., Zufall, F., & Schneider, C. (2025). Tuft cell IL-17RB restrains IL-25 bioavailability and reveals context-dependent ILC2 hypoproliferation. *Nature Immunology*, *26*(4), 567–581. https://doi.org/10.1038/s41590-025-02104-y

Gupta, A., Wodziak, D., Tun, M., Bouley, D. M., & Lowe, A. W. (2013). Loss of Anterior Gradient 2 (Agr2) Expression Results in Hyperplasia and Defective Lineage Maturation in the Murine Stomach. *Journal of Biological Chemistry*, *288*(6), 4321–4333. https://doi.org/10.1074/jbc.M112.433086

Han, S., Fink, J., Jörg, D. J., Lee, E., Yum, M. K., Chatzeli, L., Merker, S. R., Josserand, M., Trendafilova, T., Andersson-Rolf, A., Dabrowska, C., Kim, H., Naumann, R., Lee, J.-H., Sasaki, N., Mort, R. L., Basak, O., Clevers, H., Stange, D. E., … Koo, B.-K. (2019). Defining the Identity and Dynamics of Adult Gastric Isthmus Stem Cells. *Cell Stem Cell*, *25*(3), 342-356.e7. https://doi.org/10.1016/j.stem.2019.07.008

Hildersley, K. A., McNeilly, T. N., Gillan, V., Otto, T. D., Löser, S., Gerbe, F., Jay, P., Maizels, R. M., Devaney, E., & Britton, C. (2021). Tuft Cells Increase Following Ovine Intestinal Parasite Infections and Define Evolutionarily Conserved and Divergent Responses. *Frontiers in Immunology*, *12*. https://doi.org/10.3389/fimmu.2021.781108

Hong, F., Wang, X., Zhong, N., Zhang, Z., Lin, S., Zhang, M., Li, H., Liu, Y., Wang, Y., Zhao, L., Yang, X., Zhou, H., Liang, H., & Chen, Y.-G. (2025). The critical role of BMP signaling in gastric epithelial cell differentiation revealed by organoids. *Cell Regeneration*, *14*(1), 18. https://doi.org/10.1186/s13619-025-00237-x

Jin, C., Kenny, D. T., Skoog, E. C., Padra, M., Adamczyk, B., Vitizeva, V., Thorell, A., Venkatakrishnan, V., Lindén, S. K., & Karlsson, N. G. (2017). Structural Diversity of Human Gastric Mucin Glycans. *Molecular & Cellular Proteomics*, *16*(5), 743–758. https://doi.org/10.1074/mcp.M117.067983

Kalabusheva, E. P., Shtompel, A. S., Rippa, A. L., Ulianov, S. V., Razin, S. V., & Vorotelyak, E. A. (2023). A Kaleidoscope of Keratin Gene Expression and the Mosaic of Its Regulatory Mechanisms. *International Journal of Molecular Sciences*, *24*(6), 5603. https://doi.org/10.3390/ijms24065603

Kim, O., Yoon, J. H., Choi, W. S., Ashktorab, H., Smoot, D. T., Nam, S. W., Lee, J. Y., & Park, W. S. (2014). GKN2 Contributes to the Homeostasis of Gastric Mucosa by Inhibiting GKN1 Activity. *Journal of Cellular Physiology*, *229*(6), 762–771. https://doi.org/10.1002/jcp.24496

Latorre, R., Sternini, C., De Giorgio, R., & Greenwood‐Van Meerveld, B. (2016). Enteroendocrine cells: a review of their role in brain–gut communication. *Neurogastroenterology & Motility*, *28*(5), 620–630. https://doi.org/10.1111/nmo.12754

Matsuo, J., Douchi, D., Myint, K., Mon, N. N., Yamamura, A., Kohu, K., Heng, D. L., Chen, S., Mawan, N. A., Nuttonmanit, N., Li, Y., Srivastava, S., Ho, S. W. T., Lee, N. Y. S., Lee, H. K., Adachi, M., Tamura, A., Chen, J., Yang, H., … Ito, Y. (2021). Iqgap3-Ras axis drives stem cell proliferation in the stomach corpus during homoeostasis and repair. *Gut*, *70*(10), 1833–1846. https://doi.org/10.1136/gutjnl-2020-322779

Mercado-Perez, A., Hernandez, J. P., Fedyshyn, Y., Treichel, A. J., Joshi, V., Kossick, K., Betageri, K. R., Farrugia, G., Druliner, B., & Beyder, A. (2024). Piezo2 interacts with E-cadherin in specialized gastrointestinal epithelial mechanoreceptors. *Journal of General Physiology*, *156*(12). https://doi.org/10.1085/jgp.202213324

Perez White, B. E., & Getsios, S. (2014). Eph receptor and ephrin function in breast, gut, and skin epithelia. *Cell Adhesion & Migration*, *8*(4), 327–338. https://doi.org/10.4161/19336918.2014.970012

Portela-Gomes, G. M., Stridsberg, M., Johansson, H., & Grimelius, L. (1999). Co-localization of synaptophysin with different neuroendocrine hormones in the human gastrointestinal tract. *Histochemistry and Cell Biology*, *111*(1), 49–54. https://doi.org/10.1007/s004180050332

Stange, D. E., Koo, B.-K., Huch, M., Sibbel, G., Basak, O., Lyubimova, A., Kujala, P., Bartfeld, S., Koster, J., Geahlen, J. H., Peters, P. J., van Es, J. H., van de Wetering, M., Mills, J. C., & Clevers, H. (2013). Differentiated Troy+ Chief Cells Act as Reserve Stem Cells to Generate All Lineages of the Stomach Epithelium. *Cell*, *155*(2), 357–368. https://doi.org/10.1016/j.cell.2013.09.008

Tang, L., Gumkowski, F., Sengupta, D., Modlin, I., & Jamieson, J. (1996). rab3D protein is a specific marker for zymogen granules in gastric chief cells of rats and rabbits. *Gastroenterology*, *110*(3), 809–820. https://doi.org/10.1053/gast.1996.v110.pm8608891

Uhlén, M., Fagerberg, L., Hallström, B. M., Lindskog, C., Oksvold, P., Mardinoglu, A., Sivertsson, Å., Kampf, C., Sjöstedt, E., Asplund, A., Olsson, I., Edlund, K., Lundberg, E., Navani, S., Szigyarto, C. A.-K., Odeberg, J., Djureinovic, D., Takanen, J. O., Hober, S., … Pontén, F. (2015). Tissue-based map of the human proteome. *Science*, *347*(6220). https://doi.org/10.1126/science.1260419
